# Host-associated rhizobia fitness: Dependence on nitrogen, density, community complexity, and legume genotype

**DOI:** 10.1101/2020.11.20.392183

**Authors:** Liana T. Burghardt, Brendan Epstein, Michelle Hoge, Diana Trujillo, Peter Tiffin

**Affiliations:** Department of Plant and Microbial Biology, University of Minnesota, St. Paul, Minnesota; Plant Science Department, The Pennsylvania State University, University Park, Pennsylvania; Department of Plant Pathology, University of Minnesota, St. Paul, Minnesota

**Author notes:** Address correspondence to Liana T. Burghardt,. Data Accessibility:* The data supporting the results of this publication and the code necessary for these analyses will be made available via a GitHub repository after review and the raw reads have been deposited in NCBI BioProject: PRJNA401437. Authorship:* LTB designed the experiments, collected and analyzed data, and wrote the manuscript, BE designed the experiments, collected and analyzed data, MH and DT collected data, and PT designed the experiments, collected data, and wrote the manuscript.

**Keywords:** nitrogen addition, inoculation density, community complexity, *Ensifer (Sinorhizobium) meliloti*, legume-rhizobia, strain relative fitness, *Medicago truncatula*

## Abstract

The environmental context of the nitrogen-fixing mutualism between leguminous plants and rhizobial bacteria varies over space and time. Variation in resource availability, population density, and composition likely affect the ecology and evolution of rhizobia and their symbiotic interactions with hosts. We examined how host genotype, nitrogen addition, rhizobial density, and community complexity affected selection on 68 rhizobia strains in the *Ensifer meliloti* - *Medicago truncatula* mutualism. As expected, the host genotype had the most substantial effect on the size, number, and strain composition of root nodules (the symbiotic organ). The understudied environmental variable of rhizobial density had a more significant effect on strain frequency in nodules than the addition of low nitrogen levels. Higher inoculum density resulted in a nodule community that was less diverse and more beneficial but only in the context of the more selective host genotype. Higher density resulted in more diverse and less beneficial nodule communities with the less selective host. Density effects on strain composition deserve additional scrutiny as they can create eco-evolutionary feedback. Lastly, we found that relative strain rankings were stable across increasing community complexity (community complexity (2, 3, 8, or 68 strains). This unexpected result suggests that higher-order interactions between strains are rare in the context of host nodule formation and development. Taken together, our empirical work highlights the importance of developing new theoretical predictions that incorporate density dependence. Further, it has translational relevance for overcoming establishment barriers in bio-inoculants and motivating host breeding programs that maintain beneficial plant-microbe interactions across diverse agro-ecological contexts.

**IMPORTANCE:** Legume cash, forage, and cover crops establish beneficial associations with rhizobial bacteria who perform biological nitrogen fixation (BNF)—providing Nitrogen (N) fertilizer to plants without the economic and greenhouse gas emission costs of chemical N inputs. Here, for the first time, we examine the relative influence of three environmental factors that vary in agricultural fields on strain relative fitness in nodules when scores rhizobial strains compete. In addition to manipulating Nitrogen, we also use two biotic variables that have rarely been examined: the rhizobial community’s density and complexity. Taken together, our results suggest 1) breeding legume varieties that select beneficial strains despite environmental variation are possible, 2) changes in rhizobial population densities that occur routinely in agricultural fields could drive evolutionary changes in rhizobia populations, and 3) the lack of higher-order interactions between strains will allow the high-throughput assessments of rhizobia winners and losers during plant interactions.

## Introduction

Biotic interactions have important consequences for population dynamics (1), selection (2), and local adaptation (3) of interacting species. Of course, these biotic interactions do not occur in isolation and the benefits and costs of the interaction can be modified (4) by population density, the presence of additional species (5), genetics (6), and abiotic factors such as resource availability (7–10) or moisture levels (11–13). These responses (*i.e*. plasticity) can evolve if there is genetic variation in traits that modify the sensitivity of an interaction to additional environmental variables such as immunity, stress tolerance, or phenology (e.g. Ramegowda & Senthil-Kumar 2015; Garrido-Oter *et al*. 2018). Here we examine how three sources of environmental variation, resource availability, rhizobia density, and rhizobial community complexity, affect the rhizobium species (*Ensifer meliloti*) as it engages in symbiosis with its leguminous host plant (*Medicago truncatula*).

Environment dependence is a recurring theme in the study of the symbiosis between rhizobial bacteria and legume plants (16–18). In this mutualistic relationship, rhizobia convert atmospheric nitrogen (N_2_) into a plant-useable form to support host growth and reproduction while rhizobia gain carbon resources from the plant to support the growth and reproduction prior to release back into the soil (19, 20). While this relationship is commonly beneficial to the plant host (21), the magnitude of these fitness benefits to the plant depends on the identity of the rhizobial strain as well as additional environmental parameters including N, P, water- and light-availability, and temperature (17, 22–24). Experiments in which plants are inoculated with a single rhizobium strain, often at very high density, have shown that the benefits rhizobia obtain from symbiosis also can be context dependent (*e.g*. Friel & Friesen 2019; Batstone *et al*. 2020). However, in natural and agricultural populations rhizobia densities are likely to vary and multiple rhizobia strains compete for nodule occupancy and host enrichment (18). Results from single-strain experiments may not be directly translatable to multi-strain environments because between-strain competition can strongly affect nodulation success (27). Indeed, strain fitness proxies in single-strain environments are not strongly correlated with strain fitness in multi-strain communities (28, 29).

Theoretically, resource availability has the potential to shape the evolution of resource-based symbiosis (30, 31). N-availability could affect selection acting on rhizobia and plant hosts via a number of mechanisms. For instance, additional N can reduce the overall frequency of associations with hosts by reducing nodule number or size (16, 32, 33). While forming fewer nodules in high N environments may have little effect on plant fitness, it certainly reduces the chances of each rhizobia associating with a legume and, if nodules remain small, reduces the number of rhizobia released from host nodules (a major component of the fitness benefit rhizobia receive from engaging in the symbiosis). Nitrogen can also alter competitiveness among symbionts, perhaps through altering the strength of host preference (34) or enrichment via host rewards/sanctions (35, 36). Despite the appeal of theoretical predictions that additional N will reduce host dependence on rhizobia and lead to relaxed selection for rhizobial host benefit, there is limited empirical support for N-mediated shifts in competitive outcomes between rhizobia. Partner choice as measured by nodule occupancy is not strongly affected by N-addition in *Acmispon* (Regus 2014, Wendlandt *et al*. 2019) or *Medicago* (37, 38) and only weakly shifts in *Mimosa* (34). However, which strain initiates each nodule only represents the first stage of selection. Once nodules form, differential nodule growth and strain reproduction can allow some strains to increase in frequency relative to others, but again, studies suggest that N has only a limited effect on rhizobial fitness (24, 37).

Unlike N-addition, the effect of rhizobial population density on legume benefit and rhizobial fitness has received scant empirical attention. However, population densities can strongly affect the ecology and evolution of biotic interactions including hosts and pathogens (*e.g*. Schuhegger *et al*. 2006), predator and prey (*e.g*. Jaffee 2003), and plants and pollinators (*e.g*. Moeller 2004). The density of rhizobial symbionts varies over space and time. For example, in an agroecosystem, twenty years of cropping system differences resulted in four-orders-of-magnitude differences in rhizobia population densities: 6.8 × 10^6^ rhizobia gram^-1^in soy/wheat/maize rotations, 4.5 × 10^5^ in continuous soy, and 6.1 × 10^2^ in continuous maize (42). There are many reasons to suspect that selection could be density dependent; the density of rhizobia in the soil could affect the number of nodules that a host forms (43), the role of quorum sensing (44–46), and the relative importance of host-mediated vs. soil-mediated selection (47, 48). Predicting the outcome of rhizobial density on strain selection is, however, difficult. For instance, when rhizobial population densities are high more nodulation sites (root hairs) will interact with multiple strains and more nodules will be formed and thus, we might expect opportunities for plant-imposed selection on bacterial populations to increase (48). On the other hand, when rhizobial densities are high, plant control may decrease as traits that influence rhizobial competitiveness become more important (27).

Much of the empirical work on legume-rhizobia symbiosis relies on single-strain inoculations, however, in nature legume hosts often form nodules with a diverse community of strains (49, 50). Microbial community complexity can affect community assembly and interactions (51). Synthetic communities are increasingly being used to query the effect of additional community members (52–54). For example, Freidman *et al*. (2017) showed that competitive outcomes between two bacterial strains living in the guts of *C. elegans* are not affected by the presence of additional strains (56). While rhizobial strain frequency in nodules is clearly dependent on the presence of other strains (27, 37, 48), experiments have not investigated whether pairwise competitive outcomes are affected by the presence of other strains. In other words, does strain A always beat strain B regardless of which and how many other strains are present in the community?

Here we report on the extent to which strain fitness in nodules and plant traits are affected by each of three environmental factors, N-availability, population density, and the complexity of the rhizobial community to which plants are exposed (Figure 1). We measured strain communities inside host nodules using a select-and-resequence approach, a variant of evolve-and-resequence approaches (28). We inoculated plants with a community of 68 strains of the rhizobium *Ensifer meliloti* (hereafter referred to as C68). These strains were chosen to capture the majority of genetic variation present among 160 sequenced strains of *Ensifer meliloti* from a worldwide collection of strains sampled from across a broad range of environments (e.g., *Medicago* host species, geographic locals, and years (57, 58). To evaluate whether increasing the amount of N available to plants altered the strain composition of the nodule community, we grew plants with a low level of additional N (+100ml 3mMol KNO_3_ week^-1^). To evaluate the potential for rhizobia density to affect strain fitness, we inoculated plants with two rhizobial densities: low (5×10^5^ rhizobia plant^-1^) and high (5×10^7^ rhizobia plant^-1^) (Figure S1). To examine the effect of community complexity, we constructed communities of nested subsets of eight, three, and all pairwise competitions of the three. Given work showing that rhizobial fitness can strongly depend on the host genotype (28, 59–61) we examine the effect of these treatments on each of two commonly used plant genotypes. Because nitrogen addition has been widely studied, it provides context for the magnitude of less studied environmental factors.

**Figure 1:**
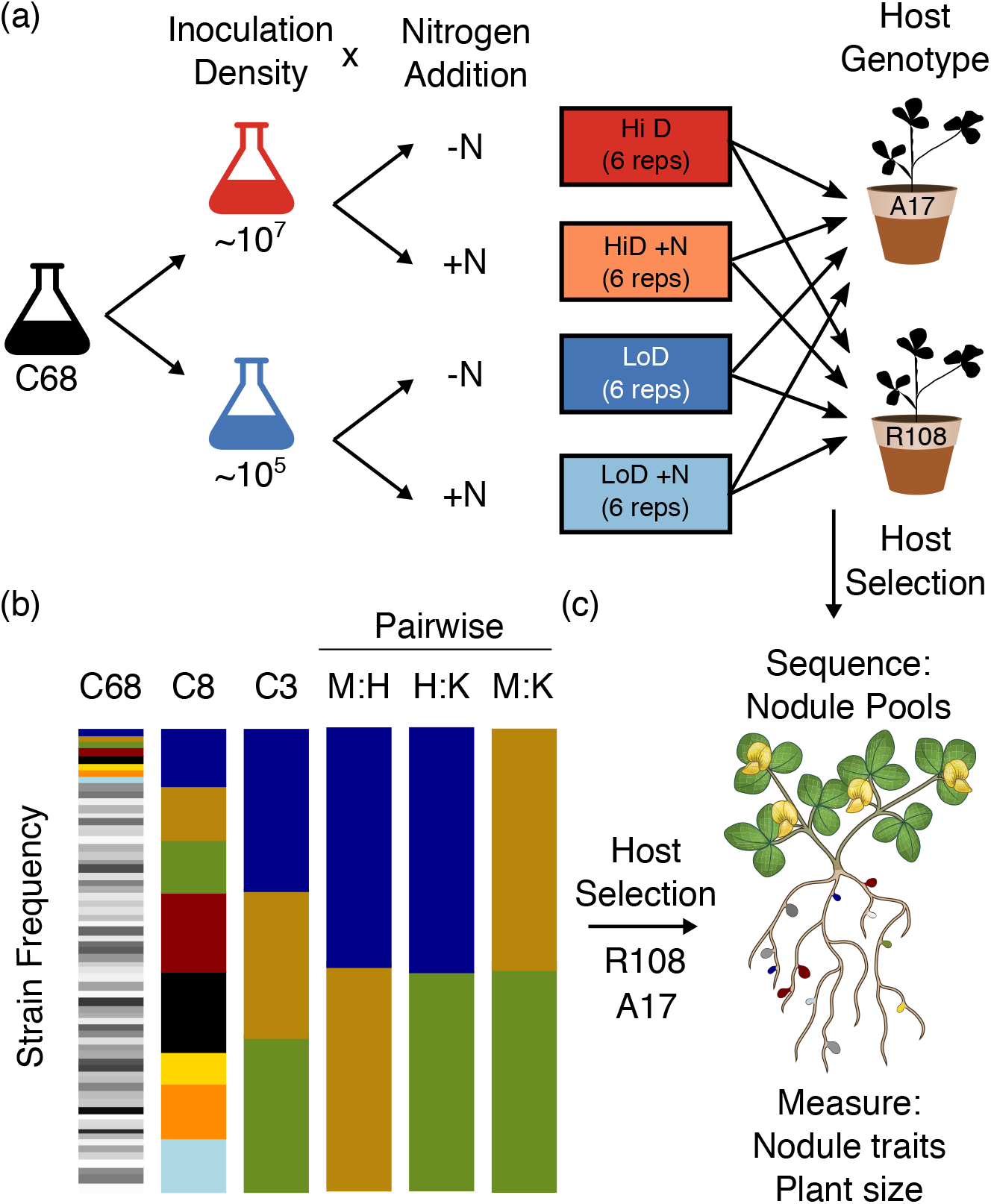
Design of the Nitrogen x Density (a) and Community complexity (b) experiments and traits measured on strains or plants in both experiments (c). All plants were harvested ~6 weeks after planting. Inoculation density is given per plant (~10 plants per pot). Colored stacked bars represent strain frequencies in each of six initial communities of 68 strains (C68-six replicate pots for each N*D treatment), eight strains (C8-five reps), three strains (C3-five reps), and two strain pairwise competitions (M:H, H:K and M:K four reps each).

Taken together, our results indicate that host-genotype has a much greater effect on strain fitness than environmental manipulations. Among the environmental manipulations, the effects of bacterial density were approximately twice as great as the effects of N-addition. Community complexity had little effect on the relative fitness rankings of rhizobial strains suggesting that higher-order interactions between strains are rare in the context of host nodule formation. Our results suggest that variation in population densities could influence ecological and evolutionary dynamics in rhizobial communities and are a factor that should be considered more explicitly in empirical, theoretical, and applied work.

## Results

### Host genotype has a large effect on strain fitness and nodule phenotypes

Consistent with previous work on the same strains (62) an overlapping set of strains (28, 61), and with an independent collection of rhizobia strains (29), the strain composition of the nodule communities was strongly affected by host genotype (Figure 2, Figure S2). Host identity had a greater effect on strain composition, Shannon’s diversity, and predicted benefit of the nodule community (Figure 2, Table 1, Table S1, Table S2), than did N-addition, inoculum density, or the complexity of the inoculum community. Relative to R108, A17 hosts produced ~ 10 times more nodules (Fig 3a) that were approximately one-tenth the size (Fig 3b) and harbored nodule strain communities that were less diverse (Fig 2b) and more beneficial (Fig 2c).

**Figure 2:**
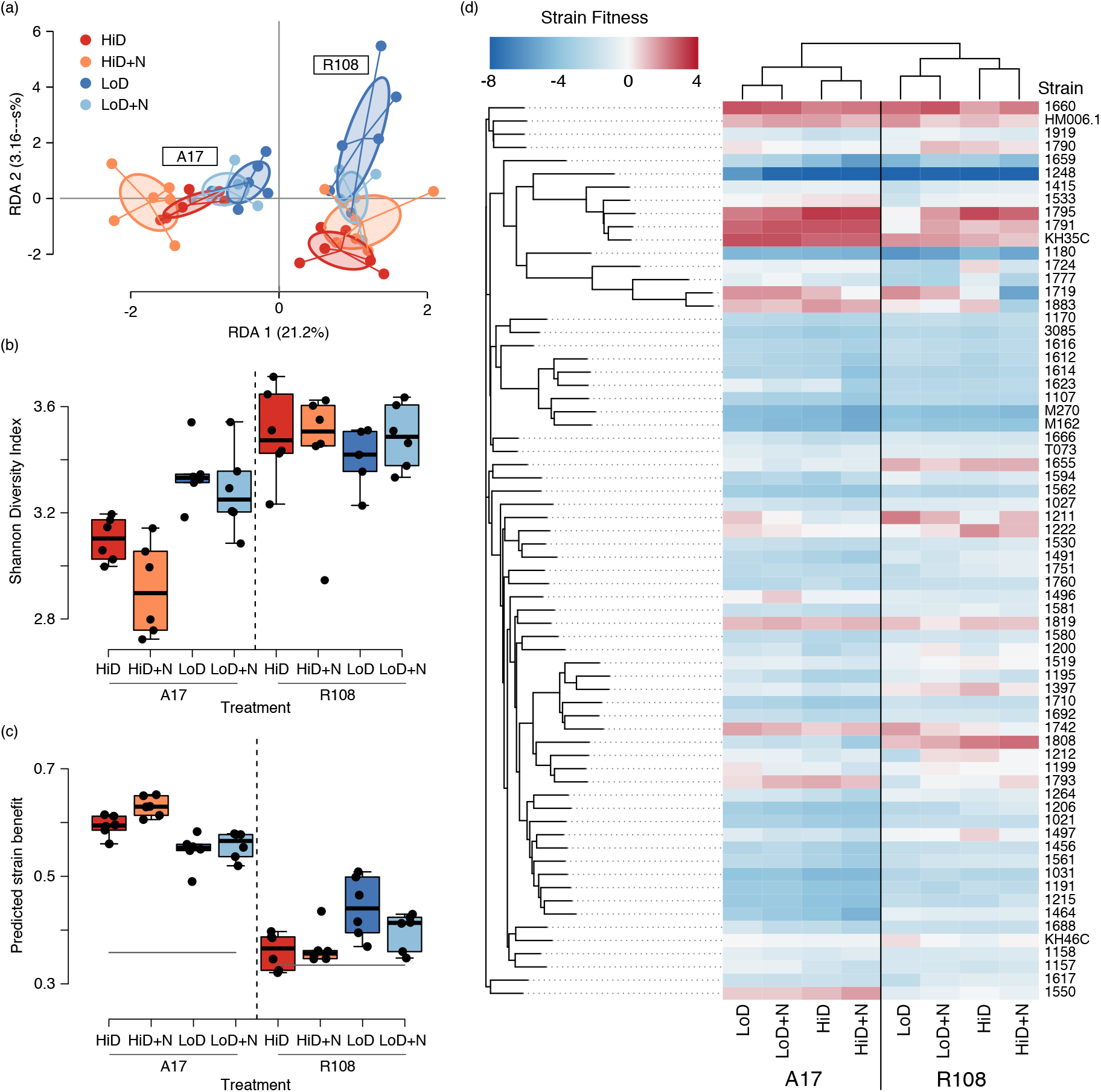
Host genotype and inoculum density affected rhizobia strain community characteristics. a) Host genotype, density, and their interaction all contributed to variation in strain composition in nodules (percent contribution is in parenthesis, Table 1, Figure S4 for RDA analysis by host genotype). b) Shannon’s diversity decreased (i.e. stronger selection) at high densities (*p*<.001) and high nitrogen (*p*=0.039) in A17. R108 nodules were always more diverse than A17 (*p*<0.001) regardless of density or nitrogen (Table S2 for full model). c) N-addition had little effect on the predicted benefit of nodule communities. However, predicted benefit significantly increased at high densities in A17 (*p*<0.001) and significantly decreased in R108 (*p*=0.003, full results in Table S3). d) Heatmap of median strain fitness values. Strains (rows) are arranged by shared SNP’s and treatments (columns) via hierarchical clustering.

**Figure 3:**
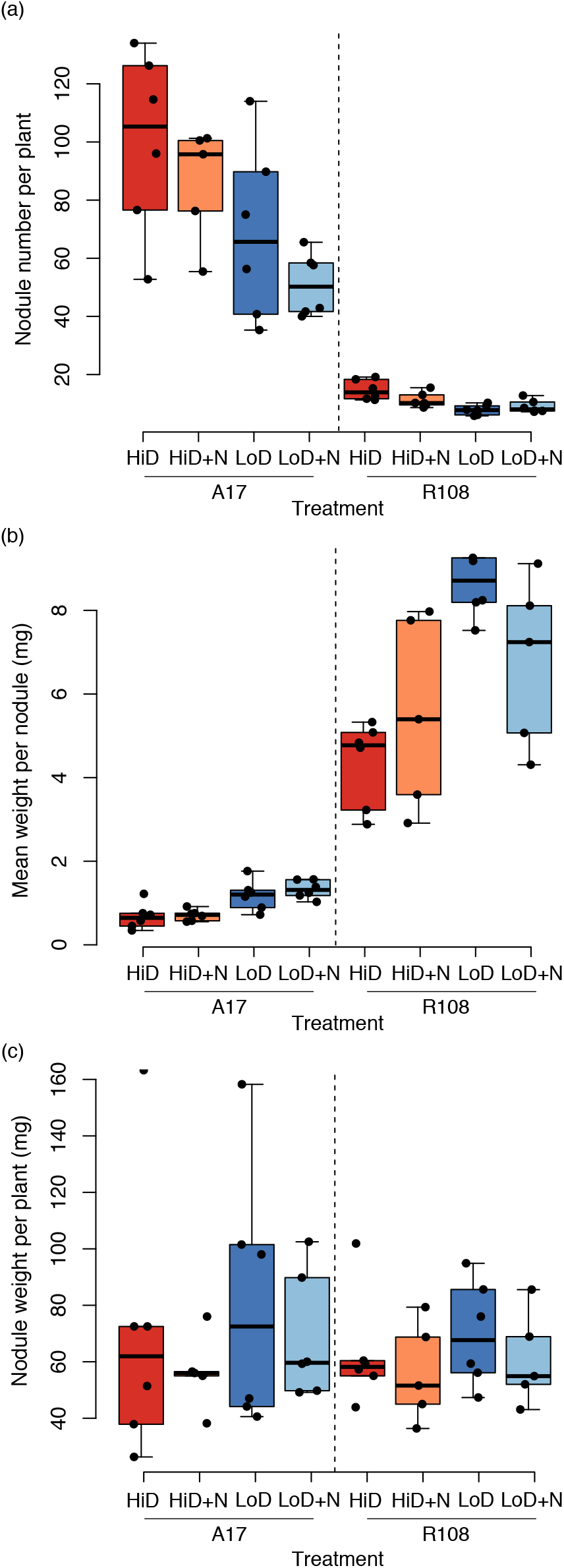
Nodule traits in Density and Nitrogen treatments. Hosts produced different numbers (a) and sizes (b) of nodules but a similar overall weight (c) across treatments. At low inoculation densities, nodule number decreased, and nodule weight increased (full results in Table S2).

**Table 1:** PERMANOVA on RDA of strain relative fitness shows effects of plant host and inoculum density are far more robust than effects of nitrogen (top). Analyses of each host separately (bottom) revealed that density and nitrogen had more significant effects with A17 than R108 hosts.

| Model and Terms | DF | Prop. Var | F | P |
| --- | --- | --- | --- | --- |
| **Both Hosts** (adj. $r^2= 0.35$) | | | | |
| Host | 1 | 0.275 | 19.81 | **< 0.001** |
| Density | 1 | 0.056 | 4.02 | **0.003** |
| Nitrogen | 1 | 0.019 | 1.35 | 0.172 |
| Host x Density | 1 | 0.038 | 2.74 | **0.015** |
| Host x Nitrogen | 1 | 0.023 | 1.67 | 0.105 |
| Density x Nitrogen | 1 | 0.018 | 1.27 | 0.223 |
| Host x Density x Nitrogen | 1 | 0.016 | 1.13 | 0.272 |
| Residual | 40 | 0.556 |  |  |
| **A17 only** (adj. $r^2= 0.20$) | | | | |
| Density | 1 | 0.217 | 6.24 | **< 0.001** |
| Nitrogen | 1 | 0.062 | 1.78 | 0.079 |
| Density x Nitrogen | 1 | 0.027 | 0.76 | 0.656 |
| Residual | 20 | 0.695 |  |  |
| **R108 only** (adj. $r^2= 0.05$) | | | | |
| Density | 1 | 0.069 | 1.68 | **0.034** |
| Nitrogen | 1 | 0.045 | 1.08 | 0.336 |
| Density x Nitrogen | 1 | 0.059 | 1.43 | 0.083 |
| Residual | 20 | 0.827 |  |  |

Note: Nitrogen was retained as a term in the sub-models because host values and variances were so divergent from each other. Proportion Variance Explained (Prop. Var); degrees of freedom (DF); P-value (P).

### N-availability weakly affected strain composition, diversity, and benefit

Because nitrogen is the primary resource rhizobia provide to their host, researchers have speculated, and modeling has shown that N-availability could alter the relative fitness of rhizobia strains. However, we found that N-availability explained only a small portion of the variance in the overall composition of the nodule communities (1.9% of the RDA variance, *p*=0.17; Figure 2), although the effect was slightly greater when hosts were analyzed separately (in A17 *p* =0.079, 6.2% of the variance and in R108 *p* = 0.336, 4.5% of the variance; Table 1, Figure S3). N-addition had a similar magnitude of effect on the diversity (Shannon’s) of the nodule community and the predicted benefit of the strain community, with the effects being greater in A17 (diversity: *p* =0.039, 8.7% of variance and strain benefit: *p* = 0.030, 7.9% of variance) than R108 (*p* =0.85, 0.17% of variance and *p* = 0.28, 3.2% of variance, respectively). See Table S1 and Table S2 for full results.

### Rhizobia inoculation density had a more substantial effect than nitrogen addition

Inoculating plants with 100-fold fewer rhizobia cells caused larger changes in community composition, strain diversity, and predicted the benefit of the nodule community than N-addition (Figure 2). Moreover, the effect of density depended on the host. With A17 hosts, density explained 21.7% (*p*=0.001) of variance in community composition vs. 6.2%(*p*=0.079) of the variance explained by N-addition (Figure S3b). With R108, density explained 6.9% (*p*=0.034) of the variance vs 4.5 % (*p*=0.336) explained by N-addition (Figure S3c). Strain fitness shifts in response to density are so strong they are clear even in an un-constrained PCA analysis of strain fitness (Figure S4). Density affected the diversity of the nodule community in opposite directions in the two hosts. Compared to the low-density inoculation, the high-density inoculation resulted in A17 nodule communities being less diverse (based on Shannon’s diversity) and more beneficial (both *p* < 0.001). By contrast, with R108, the low-density inoculation resulted in a less beneficial nodule community (*p* < 0.003, Table S1, S2).

### Relative fitness rankings are similar across communities of increasing complexity

In nature, the complexity of rhizobial communities varies in both space and time. We found that although the absolute frequency of each of the three focal strains was strongly affected by the presence of additional strains in the inoculum, we found little evidence that additional strains in the inoculum community altered strain rankings relative to each other. In other words, with minor exceptions, the more frequent strain in pairwise competitions was also the more frequent strain in more complex communities (Table 2). The minor exceptions involved two strains (KH46c and HM006.1) that had nearly equal frequencies in pairwise inoculations and nearly equal frequency when part of more strain-rich communities (Fig S5, S6; Table 2). Similarly, the relative-frequency ranking of strains when plants were inoculated with three strains (C3) were almost always the same as the relative ranking of those strains when they were part of a 68-strain inoculum community (Figure 4a-c) and the relative ranking of the strains in C8 treatment was nearly always the same as in the C68 treatment (Figure 4d-f). These results suggest that strain competitiveness for nodulation formation is relatively robust to the presence of additional strains, and strains with higher frequency in pairwise competitions will also have higher frequencies in more complex communities. Interestingly, we did observe hints of negatively frequency dependence – extremely low-frequency strains tended to be more frequent in more complex communities (e.g. M162 in C3 vs. C68 and T073 in C8 vs. C68 Fig 4c,f).

**Figure 4:**
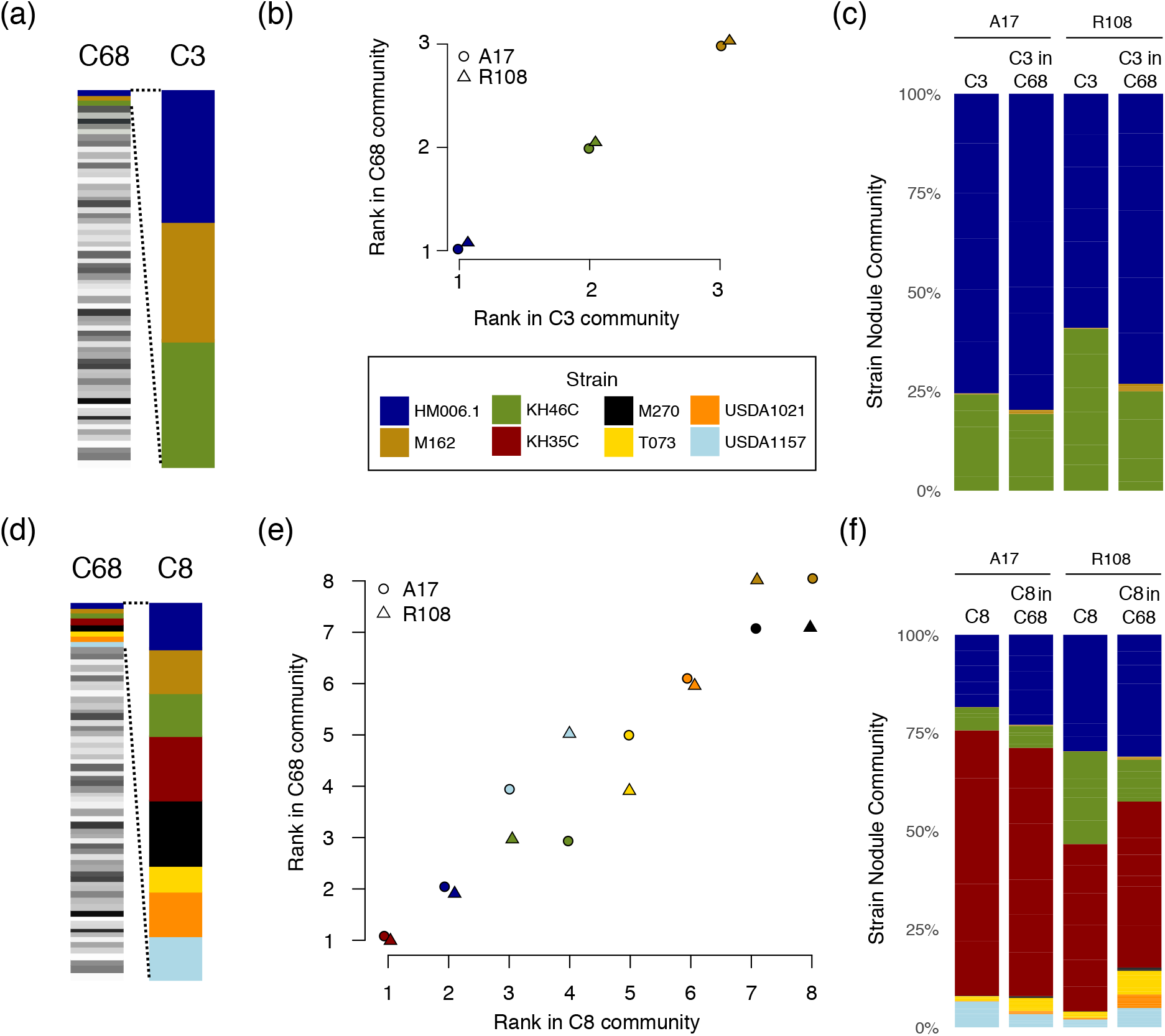
Rank order of relative strain frequencies in the nodule communities were identical when plants were inoculated with either the 3-strain (C3) or 68-strain (C68) community (a-c), and highly similar when inoculated with either the 8-strain (C8) or C68 community (d-f). (a,d) show nested strain subsets of the initial communities. (b, e) Strain ranks as determined based on mean strain frequency across five (C3 and C8) or six (C68) replicates for A17 (triangles) and R108 (circles) and depicted in (c, f). In A17, the third and fourth-ranked strains USDA1157 and KH46C swapped places, and in R108, T073 and USDA1157 swapped and the rarest strains M270 and M162 swapped. See Fig S5 for additional contrasts and Fig S6 for variation between individual replicates.

**Table 2:**
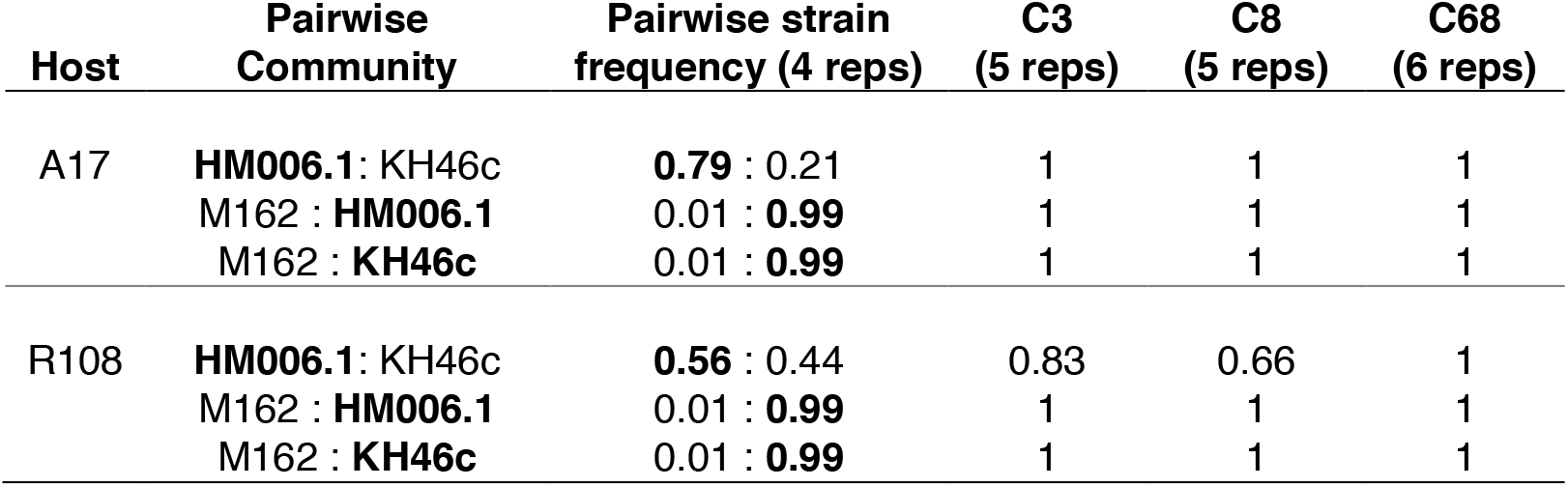
Rankings of strain frequencies in two strain communities were consistant across increasingly complex communities: three strains (C3), eight strains (C8), and 68 strains (C68). The higher frequency strain is shown in bold followed by the proportion of replicates in which the higher frequency strain also had higher relative frequency in the more complex communites.

| Host | Pairwise Community | Pairwise strain frequency (4 reps) | C3 (5 reps) | C8 (5 reps) | C68 (6 reps) |
| --- | --- | --- | --- | --- | --- |
| A17 | **HM006.1**: KH46c | **0.79** : 0.21 | 1 | 1 | 1 |
|  | M162 : **HM006.1** | 0.01 : **0.99** | 1 | 1 | 1 |
|  | M162 : **KH46c** | 0.01 : **0.99** | 1 | 1 | 1 |
| R108 | **HM006.1**: KH46c | **0.56** : 0.44 | 0.83 | 0.66 | 1 |
|  | M162 : **HM006.1** | 0.01 : **0.99** | 1 | 1 | 1 |
|  | M162 : **KH46c** | 0.01 : **0.99** | 1 | 1 | 1 |

### Effects of N, D, and complexity on nodule and plant traits were host specific

The effect of N-addition and inoculum density on plant phenotypes were host-specific (Table S3). In A17, increased rhizobial density resulted in plants forming more nodules (*p*=0.004; Fig 3a) that weighed less (*p*<0.001; Fig 3b). In R108, an increase in rhizobia density resulted in plants forming more, smaller nodules, but only in the N-addition environment (Nitrogen x Density *p* = 0.036 and *p* = 0.033, respectively). Neither N-addition nor rhizobia density had measurable effects on either above or belowground biomass. However, both N (*p*=0.029) and to a lesser extent density (*p*=0.087) decreased the root to shoot ratio in A17 but not in R108 (Fig S7). The complexity of the inoculum community had little effect on nodule number or vegetative biomass (Fig S8; Table S4), although it did have minor effects on nodule weight in A17 but not R108 (Host x Complexity *p*= 0.018), and root weight (*p* = 0.053) of both hosts.

## Discussion

The symbiosis between legumes and rhizobia plays an important ecological role by contributing nitrogen to natural and agricultural ecosystems (63–65). Here we asked how host genotype and each of three environmental factors likely to vary spatially and temporally affect the rhizobia community in nodules. Despite being the key benefit that rhizobia provide plants, Nitrogen had relatively minor effects on the strain composition, diversity, and beneficialness of nodule communities. Similarly, the complexity of the inoculum community (number of strains used to inoculate a plant) had only minor effects on relative strain rankings, suggesting that higher-order interactions among strains do not alter the pairwise rankings of strain success in the context of nodule formation. Interestingly, inoculum density had large and host-specific effects on the nodule community—density had a more significant effect on strain composition, diversity, and benefit in the host (A17) that imposes greater selection on the rhizobia population.

Theoretically, the availability of alternative sources of the symbiotic resource can alter selection on beneficial symbionts (30, 31, 66). When mutualistic relationships are considered in a market framework, external N improves the bargaining power of the host and increases the relative fitness of less beneficial strains (31). Similarly, relaxed selection for symbiont benefit could occur because it is less costly for the host to allocate resources to a less beneficial partner if it can obtain the resource from an alternative source (30). Despite the potential, we found that the addition of small amounts of N had little effect on rhizobial strain composition, diversity, and the host benefits of the nodule community. Indeed, in one host genotype (A17), selection for beneficial strains was stronger, not weaker, in the N-addition environment. Our findings of subtle effects of N-addition are consistent with several other studies that have examined the composition of strains that form nodules (37, 38, 67–69). Similarly, our results align with two studies that assessed composite strain fitness across all nodules formed by a plant. Nitrogen addition did not affect rhizobial composition in pools of 4-week-old *Medicago* nodules inoculated with a mixture of two *Ensifer* species or a mixture of Fix- and Fix+ strains (37) and had weak effects on strain fitness two species of *Acacia*—drought and phosphorous addition treatments had larger effects (24).

Although our results are consistent with these short-term (single growing season) studies, it is clear the addition of N over longer periods of time can affect the symbiosis. Many legumes regulate nodule formation and symbiotic nitrogen fixation when high levels of abiotic N are available, presumably reflecting a cost of forming and maintaining the symbiosis—(33). For instance, Weese *et al*. found rhizobia in a grassland community to be less beneficial to hosts after 22 years of N addition (70, 71). One way to reconcile discrepancies between short and long-term experiments is to note that other parameters that influence absolute fitness could change even if relative fitness is not altered by N-addition. For instance, reductions in host population sizes, nodule numbers, or nodule size could all increase the importance of rhizobial adaptation to environments external to the host and result in a population of strains with reduced host benefit via selective tradeoffs or drift (18, 28, 72–74). To test such a hypothesis will require follow-up experiments that hold relative frequencies of strains, i.e., relative fitness, constant while varying rhizobial population densities and frequency of exposure to host and non-host environments.

Whereas the effects of N on the legume-rhizobia symbiosis have been the focus of many studies, the potential for density-dependent selection shaping legume-rhizobia symbiosis has received scant attention. Nevertheless, rhizobial density varies widely in nature (e.g., 10^2^ −10^6^ per gram; Yan *et al*. 2014). The 100-fold difference in inoculum density we examined resulted in host-dependent shifts in the composition of the nodule community, with increased density resulting in more beneficial strains being favored in one host and less beneficial strains being favored in the other host. The results raise questions about the generality of laboratory experiments that rely on extremely high densities of rhizobia—presumably to ensure that hosts will not be rhizobia limited. Interestingly, while nodules sampled from field-grown plants are nearly always occupied by a single rhizobia strain (69, 75, 76), the probability of one nodule being infected by multiple strains increases with rhizobial density (47, 48). Whether strain competition occurs via direct interactions within a nodule vs. competition between nodules inhabited by different strains can affect mechanisms of rewards/sanction and the scale at which selection occurs (77).

Although most experiments are conducted with relatively simple rhizobia communities, in nature, legume hosts are exposed to many strains (42, 50), as well as other microbes that might affect the establishment of the symbiosis (23, 78). Studies of species-level interactions between microbes show that adding an additional species can sometimes modify interactions between two species and even cause transitions from mutualisms to antagonisms (79). We did not find evidence for these sorts of shifts among *E. meliloti* strains. The rank order relationships among focal strains were mainly unchanged when we added additional strains. Because we know of no other studies of mutualism that examine intra-specific bacterial variation, it is not easy to contextualize these results. Still, the apparent lack of shifts in pairwise interactions indicates that mixed-strain experiments could help classify strain interactions more broadly. For instance, a similar methodology could be used to assess competitive outcomes between scores of mutant strains and a wildtype strain in a single experiment instead of conducting scores of pairwise competition experiments that preclude direct comparisons between mutants. Similarly, our results suggest that multi-strain inoculation experiments could be used in breeding pipelines to screen for rhizobial strains with high fitness on specific host varieties.

Experimental evolution studies, such as this one, allow assessment of the relative importance of deterministic and stochastic processes in species interactions (80–82). The two host genotypes differed markedly in the characteristics of nodules in ways that could influence the evolution of rhizobial symbionts. For instance, A17 made many small nodules while R108 made a few large nodules, a pattern found in all nitrogen, density, and inoculum environments. Despite these differences, overall investment in nodule tissue per plant was very similar for both hosts. Decreases in nodule numbers at low inoculum densities were offset by increases in nodule size (a pattern previously observed by Singleton & Tavares 1986). While rarely discussed from the rhizobia perspective, these striking genotypic differences could have significant consequences for the evolution of rhizobial symbionts. Our observation of low nodule numbers and low selectivity of R108 host plants provides a testable hypothesis for the increased variability in strain fitness amongst replicates. Increased stochasticity in strain fitness outcomes even when pooling nodules from ten or more host plants could influence rhizobial evolution— not by directly determining rhizobial fitness—but by increasing the strength of drift. Our results add to the increasing number of studies documenting the potential importance of stochastic processes (84–86) in host-microbe interaction and hint that genetically-influenced host traits can drive differences in the relative importance of these processes. Screening additional accessions of *Medicago truncatula* for these traits and expanding the scale of genetic variation to individual host genes and multiple Medicago species will provide a broader understanding of the range of host genetic control on symbiont evolution.

### Conclusions

Using a high-throughput methodology, we evaluated the context-dependence of host-dependent strain fitness in legume-rhizobia symbiosis. Our finding that host genetic variation is a consistent driver of rhizobial relative fitness across environmental perturbations suggests it is possible to identify the genomic basis of strain x host interactions (87–89) and use that information to identify successful rhizobial strains even when environmental conditions change across years or locations. Our results also suggest that the selection hosts impose on rhizobial populations might depend on the density of the rhizobia population in the soil. From a translational perspective, understanding density-dependent selection could aid in developing beneficial inoculants that overcome establishment barriers (90).

## Material and methods

### Constructing Communities

To form the rhizobia communities, we grew each of 68 strains in 3ml Tryptone yeast media (6g tryptone, 3g yeast extract, 0.38g CaCl_2_ per L) for three days and then combined an equal volume of each culture to generate a community (C68) with approximately equal representation of each strain (median strain frequency 0.014, range 0.009-0.02). We used the same method to form the eight (C8), three (C3), and three two-strain communities (M:H, M:K, and H:K). All strain names are listed in Fig 2d.

### Experimental Details

Seeds of two host genotypes A17 var. Varma and R108 (Medicago HapMap accession numbers HM101 and HM340, Stanton-Geddes et al. 2013) were bleached, rinsed, scarified with a razor blade, stratified on wet filter paper at 4°C in the dark for two days, and then allowed to germinate at room temperature for one day. We chose these genotypes because they 1) are commonly used for molecular genetics work and 2) differ in nodule strain communities, transcriptomes, and symbiotic traits. Twelve germinated seeds were then planted in each of 45 1L pots filled with sterilized Sunshine Mix. When seedlings were three days old, 100 ul of each rhizobial community diluted in 9.9 ml 0.85% NaCl w/v solution were used to inoculate each pot (approximately 10^8^ cells, except for the low-density treatment, which was inoculated with ~ 10^6^ cells, Fig S1). Plants were fertilized with 150ml of N-free fertilizer (Bucciarelli *et al*. 2006 see Burghardt et. al. 2018 for details) once a week and watered with sterile water as needed. Six weeks after planting, we sampled ~300-500 (A17) or ~100-200 (R108) nodules from the plants in each pot (Fig S9). Because genotype A17 produces 5-10 times more nodules than genotype R108, this sample represents nodules from all 10-12 plants in pots with R108 and ~ 6 plants from A17 pots. This sample size is designed to ensure we have statistical power to overcome stochastic processes underlying nodule formation, given the 68 potential strain partners. Nodule pools were crushed, and we used a series of differential centrifugation steps to enrich for and pellet undifferentiated bacteria (Burghardt et al. 2018). Pellets were stored at −20°C until we extracted DNA using the UltraClean Microbial DNA Isolation Kit (no. 12224; Mo Bio Laboratories). In addition to harvesting the nodules, we measured on a per plant basis: nodule number, nodule fresh weight, and vegetative and root biomass (dried at 60°C for 72 hours). For each host genotype, we sampled six nodule pools for the four Nitrogen x Density C68 treatments, five nodule pools of the C8 and C3 treatments, and four nodule pools for each of the three pairwise community treatments.

### Strain frequencies

We estimated the frequency of each strain in each nodule pool using the method in Burghardt et al. 2018. In brief, DNA isolated from each replicate was sequenced on an Illumina HiSeq 2500 (NexteraXT libraries, 125 bp paired-end reads, 3.6-9.4 million read pairs library^-1^). Reads were trimmed with TrimGalore! (v0.4.1) using default settings, except with minimum read length = 100 bp, quality threshold = 30, and minimum adapter match = 3. We used bwa mem (v0.7.17; Li and Durbin, 2010) with default settings to align reads to the *E. meliloti* USDA1106 genome (Nelson *et al*. 2018; NCBI BioProject: PRJNA388336). Using prior genome sequencing data on the 68 strains, we identified SNPs segregating in each of the sequenced communities using FreeBayes (v1.2.0-2-g29c4002; (93) with a minimum read mapping quality of 30. After cleaning and alignment, the median read depth per sample was 65X (range 29X-115X). To estimate strain frequency, we used only SNPs for which every strain had an unambiguous call. We then estimated the frequency of each strain in each sample using HARP (94) as described in Burghardt *et al*. (2018). Briefly, this method estimates the likelihood that each read comes from each strain and summarizes this signal across all reads to estimate strain frequencies.

### Nodule community measurements

Based on strain frequencies, we calculated three nodule community metrics for each replicate pot: composition, diversity, and host benefit. We estimated community composition as the fold change in the frequency of each strain (*q_x_*) in a nodule community relative to the mean frequency of that strain across four sequencing replicates of the initial community 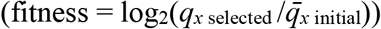. This transformation normalizes the frequency distribution and controls for small differences in initial strain frequency (median *q*_*x* initial_=0.0092, 5%-95% quantile: 0.0054-0.0154). We estimated community diversity as the exponent of Shannon diversity, which we calculated using the ‘renyi’ function in the vegan package of R. We calculated predicted host benefit as the sum of the per strain frequency in nodules multiplied by the dry plant weight from a single-strain inoculation experiment involving both A17 and R108 (data from (Burghardt *et al*. 2018):

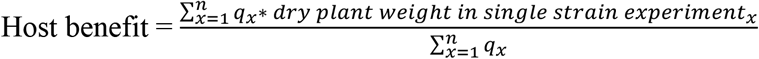

We scaled each host-specific dataset from zero (full occupancy by the least beneficial strain) to one (full occupancy by the most beneficial strain). We omitted strains for which we did not have single-strain plant weight estimates (9 strains for A17 and 29 strains for R108). While missing strains may reduce the power to make between host comparisons, missing strains span the phylogeny, and there is no reason to suspect that our subsamples are biased.

### Statistical analysis of community measures

We used redundancy analysis (RDA, “rda” in the vegan R package; Oskanen et al. 2017) and ANOVA to analyze the effects of Host genotype (H), Density (D), and Nitrogen (N) and their interactions on each of the nodule community measures. To collapse the dimensionality of the strain relative fitness data and analyze shifts in relative fitness across treatments, we used RDA. RDA fits a multivariate linear regression to centered and scaled data and then uses principal component analysis (PCA) to decompose the major axes of variation in the fitted parameters. The adjusted R^2^ of each RDA model provides an estimate of the proportion of variance in relative fitness explained by the model predictor(s). We permuted the data to determine the probability that fitness differences occurred by chance (‘anova’ function, 999 permutations). We used an ANOVA (‘lm’ and ‘anova’) to analyze strain diversity to test for differences amongst treatments. To analyze host benefit, we used an ANOVA (‘lm’ and ‘anova’). Because the effects of D and N were host-dependent (H x D, H x N interactions), we also analyzed the effects of N, D, and their interaction for each host separately.

### Analysis of rank order and community complexity

For each pairwise competition of the three strains used to form a three-strain community (C3), we identified the strain with higher strain frequency and asked if the same strain was at higher frequency in the more complex three strain community. We also examined whether strain frequency rankings in the C3 community remained the same in the eight- (C8) and sixty-eight- (C68) strain communities. Likewise, we examined whether the frequency rankings of each of the strains in the C8 community remained the same in the C68 community.

### Statistical analysis of plant phenotypes

We used an ANOVA (‘lm’ and ‘anova’) to test for the effects of host genotype, inoculum density, and nitrogen level and their interactions on six plant traits (nodule number, average nodule weight, nodule weight per plant, shoot biomass, root biomass, and root to shoot ratio). The first two traits violated the assumption of homogeneity of variance between hosts, so we focus on host-specific analyses. To evaluate the effect of community complexity on plant phenotypes, we ran a model with Host (H) and community complexity (C; 2, 3, 8, and 68 strains) and their interaction as continuous predictors. Although we test for the among-treatment differences in plant phenotypes, the experiments were designed to evaluate the effect of treatments on strain fitness. They were not well powered to detect among-treatment differences in plant phenotypes.

## Acknowledgments

We thank Roxanne Denny for help with seed sources. Computational resources were provided by the Minnesota Supercomputing Institute (MSI) at the University of Minnesota. This work was supported by the National Science Foundation (NSF) awards IOS-1724993 and IOS-1856744 to LTB and PT. Any opinions, findings, conclusions, or recommendations expressed in this material are those of the authors and do not necessarily reflect the views of the NSF. LTB’s work was additionally supported by the USDA National Institute of Food and Agriculture Federal Appropriations under PEN04760 and Accession #1025611.

## Notes

### Competing Interest Statement

The authors have declared no competing interest.

