## Supplemental Figures and Tables for "Host-associated rhizobia fitness: Dependence on nitrogen, density, community complexity, and legume genotype"

Supplemental Table S1: Shannon diversity in nodules is influenced by inoculum density (Density) and nitrogen addition (Nitrogen) in A17 but not R108. Full ANOVA model with both hosts (top) and separate models for each host (bottom). For each term, we show degrees of freedom (df), model sum of squares (Sum Sq.) and *p* value. For the full model, we also include F-values.

| Bost Hosts | Df | Sum Sq. | F value | *p* |
|---|---|---|---|---|
| Host | 1 | 1.07 | 44.0 | **<0.001** |
| Density | 1 | 0.25 | 10.1 | **0.003** |
| Nitrogen | 1 | 0.037 | 1.5 | 0.23 |
| Host x Density | 1 | 0.32 | 13.1 | **0.001** |
| Host x Nitrogen | 1 | 0.057 | 2.3 | 0.14 |
| Density x Nitrogen | 1 | 0.052 | 2.1 | 0.15 |
| Host x Density x Nitrogen | 1 | 0.000 | 0.002 | 0.97 |
| Residuals | 40 | 0.97 | | |

| | | A17 | | R108 | |
|---|---|---|---|---|---|
| | Df | Sum Sq. | *p* | Sum Sq. | *p* |
| Density | 1 | 0.56 | **<0.001** | 0.0024 | 0.78 |
| Nitrogen | 1 | 0.092 | **0.039** | 0.0011 | 0.85 |
| Density x Nitrogen | 1 | 0.024 | 0.27 | 0.027 | 0.35 |
| Residuals | 20 | 0.38 | | 0.60 | |

Supplemental Table S2: Predicted host benefit in nodules is strongly affected by inoculum Density with both A17 and R108 hosts, whereas the effect of Nitrogen addition is greater in A17 that R108. Full ANOVA model with both hosts (top) and separate models for each host (bottom).

| Bost Hosts | Df | Sum Sq. | F value | *p* |
|---|---|---|---|---|
| Host | 1 | 0.45 | 382 | **<0.001** |
| Density | 1 | 0 | 0.02 | 0.88 |
| Nitrogen | 1 | 0 | 0.05 | 0.82 |
| Host x Density | 1 | 0.041 | 35.2 | **<0.001** |
| Host x Nitrogen | 1 | 0.005 | 4.58 | **0.038** |
| Density x Nitrogen | 1 | 0.005 | 3.96 | 0.054 |
| Host x Density x Nitrogen | 1 | 0 | 0.38 | 0.54 |
| Residuals | 40 | 0.047 | | |

| | | A17 | | R108 | |
|---|---|---|---|---|---|
| | Df | Sum Sq. | *P* | Sum Sq | *p* |
| Density | 1 | 0.022 | **<0.001** | 0.02 | **0.003** |
| Nitrogen | 1 | 0.003 | **0.03** | 0.002 | 0.28 |
| Density x Nitrogen | 1 | 0.001 | 0.19 | 0.004 | 0.147 |
| Residuals | 20 | 0.012 | | 0.035 | |

Supplemental Table S3: The influence of nitrogen addition (Nitrogen), inoculum density (Density), and host genotype (Host) on nodule and plant traits (Trait~Host*Density*Nitrogen). For each term, we show model sum of squares and *P* value category (****p* <0.001, ** *p* <0.01, * *p* <.05, • *p* < 0.1). For the three traits with significant interactions in a full ANOVA model we ran sub models for each host separately (bottom). There 39 residual degrees of freedom for each full model and 19 for each sub-model.

| Terms | Nodule Number per plant | Weight per Nodule | Nodule weight per plant | Veg. weight per plant | Root weight per plant | Root: Shoot |
|---|---|---|---|---|---|---|
| Host | **50030***** | **340***** | 547 | **299979***** | **453088***** | 0.1 |
| Density | **4169**** | **36***** | 801 | 15823 | 891 | 0.1 |
| Nitrogen | *889•* | 0.4 | 1302 | 18764 | 2407 | 0.2 |
| Host:Density | **2451**** | **18**** | 83 | 5566 | 25602 | *0.4•* |
| Host:Nitrogen | 639 | 1 | 96 | 14848 | 13797 | **0.6**** |
| Density:Nitrogen | 2 | **7*** | 1 | 461 | 1115 | 0 |
| Host:Density:Nitrogen | 47 | **8*** | 8 | 1808 | 10561 | 0.1 |
| Residuals | 11725 | 53 | 32078 | 292074 | 389082 | 4.1 |

| Terms | Nodule Number per plant | | Weight per Nodule | | Root:Shoot | |
|---|---|---|---|---|---|---|
| | A17 | R108 | A17 | R108 | A17 | R108 |
| Density | **6563**** | **123***** | **2***** | **52***** | *0.4•* | 0.1 |
| Nitrogen | 1454 | 8 | 0 | 1 | **0.8**** | 0 |
| Density:Nitrogen | 16 | **33*** | 0 | **15*** | 0 | 0.1 |
| Residuals | 11594 | | 1.5 | 53 | 2.7 | 1.4 |

Supplemental Table S4: ‘root weight per plant’ weight per nodule’ increase with community complexity in R108 but not A17 hosts. In contrast, ‘nodule weight per plant’ was not affected by host genotype or community complexity. Top) Results from a full model (trait~host+complexity+ host*complexity). Bottom) Results from sub-models for each host for ‘weight per nodule’ and ‘root weight per plant’. For each term, we show model sum of squares and $p$ value category (*** $p$<0.001, ** $p$ <0.01, * $p$ <.05, • $p$ < 0.1)

| Terms | DF | Nodule Number per plant | Weight per Nodule | Nodule Weight per plant | Vegetative weight per plant | Root weight per plant | Root: Shoot ratio |
|---|---|---|---|---|---|---|---|
| Host | 1 | **76295*** ** | **117.9*** ** | 0.000 | **0.303*** ** | **0.435*** ** | 0.003 |
| Complexity | 1 | 409 | **3.06*** | 0.001 | 0.009 | *0.039•* | 0.105 |
| Host:Complexity | 1 | 856 | **3.14*** | 0.000 | 0.000 | 0.002 | 0.000 |
| Residual | 56 | 17443 | 29.7 | 0.028 | 0.227 | 0.548 | 6.413 |

| Terms | | Weight per Nodule | | Root weight per plant | |
|---|---|---|---|---|---|
| | DF | A17 | R108 | A17 | R108 |
| Complexity | 1 | 0.00024 | **6.2006*** | .1205 | **0.0277*** |
| Residual | 28 | 1.34 | 53 | .3769 | 0.1713 |

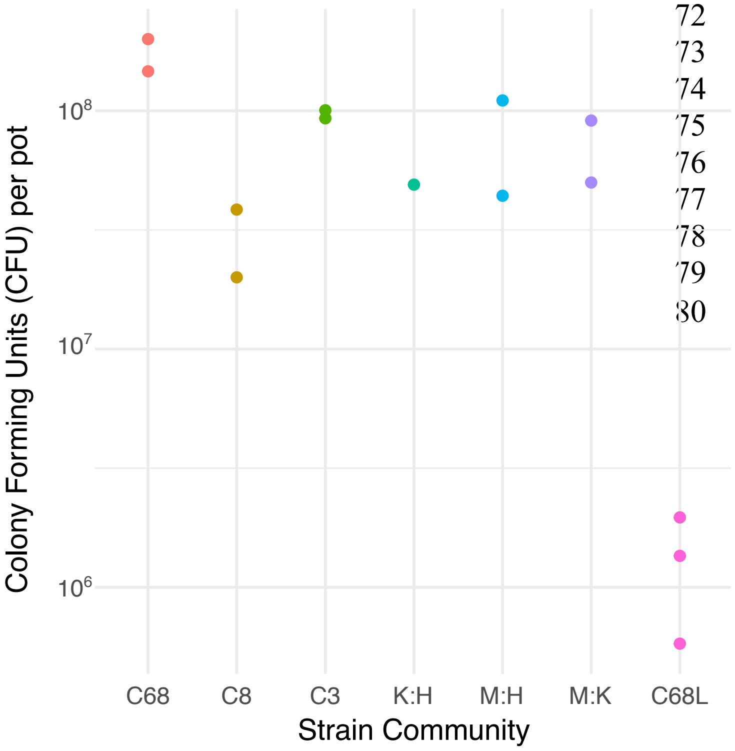


Supplemental Figure S1: Densities of initial inoculum communities applied per pot as measured by dilution plating and counts of colony-forming units. Since each pot contained ~10-12 plants, each plant received ~$10^7$ cells in high-density treatments and ~$10^5$ cells in the low-density treatment.

Supplemental Figure S2: Community composition summarized across replicates for each Host*Nitrogen*Density treatment. Strains colors designate the focal strains used in the reduced community complexity experiment.

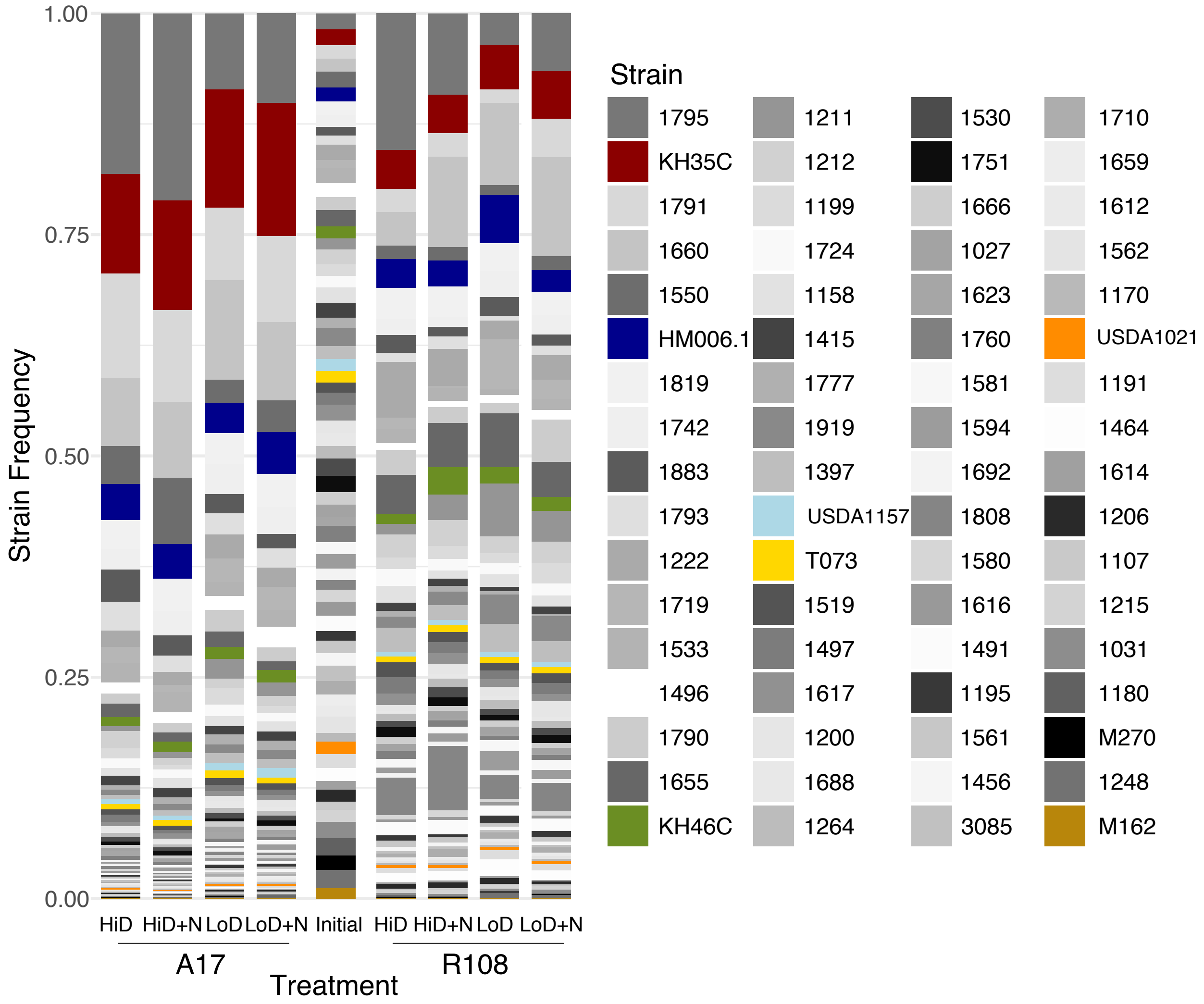

Figure S3: Visualization of the first two axes from an RDA analysis of the influence of inoculum density and nitrogen addition on strain relative fitness separated by host. Density and Nitrogen have considerably larger effects on the strain composition of A17 nodules (a) than R108 (b) nodules (full statistical results in Table 2).

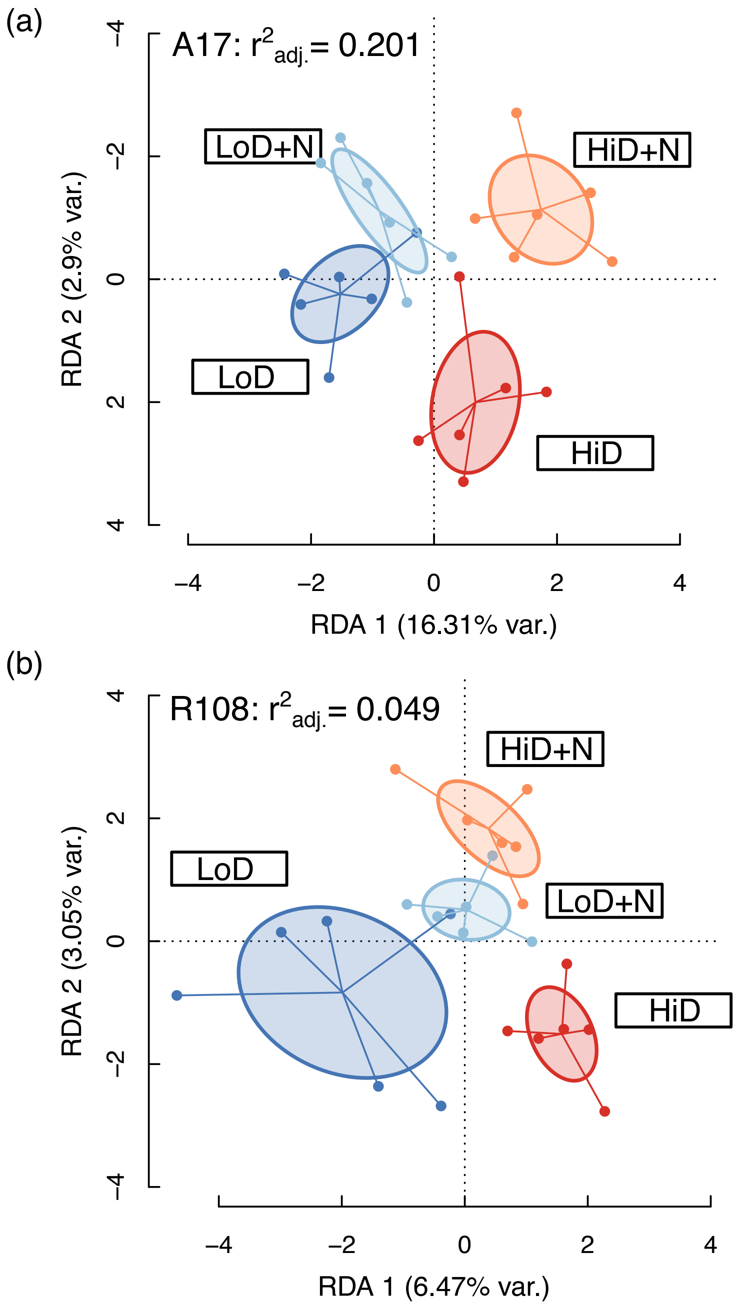

Supplemental Figure S4: Principal Component Analysis (PCA) of strain relative fitness combined across hosts (a) and for A17(b) and R108 (c) alone.

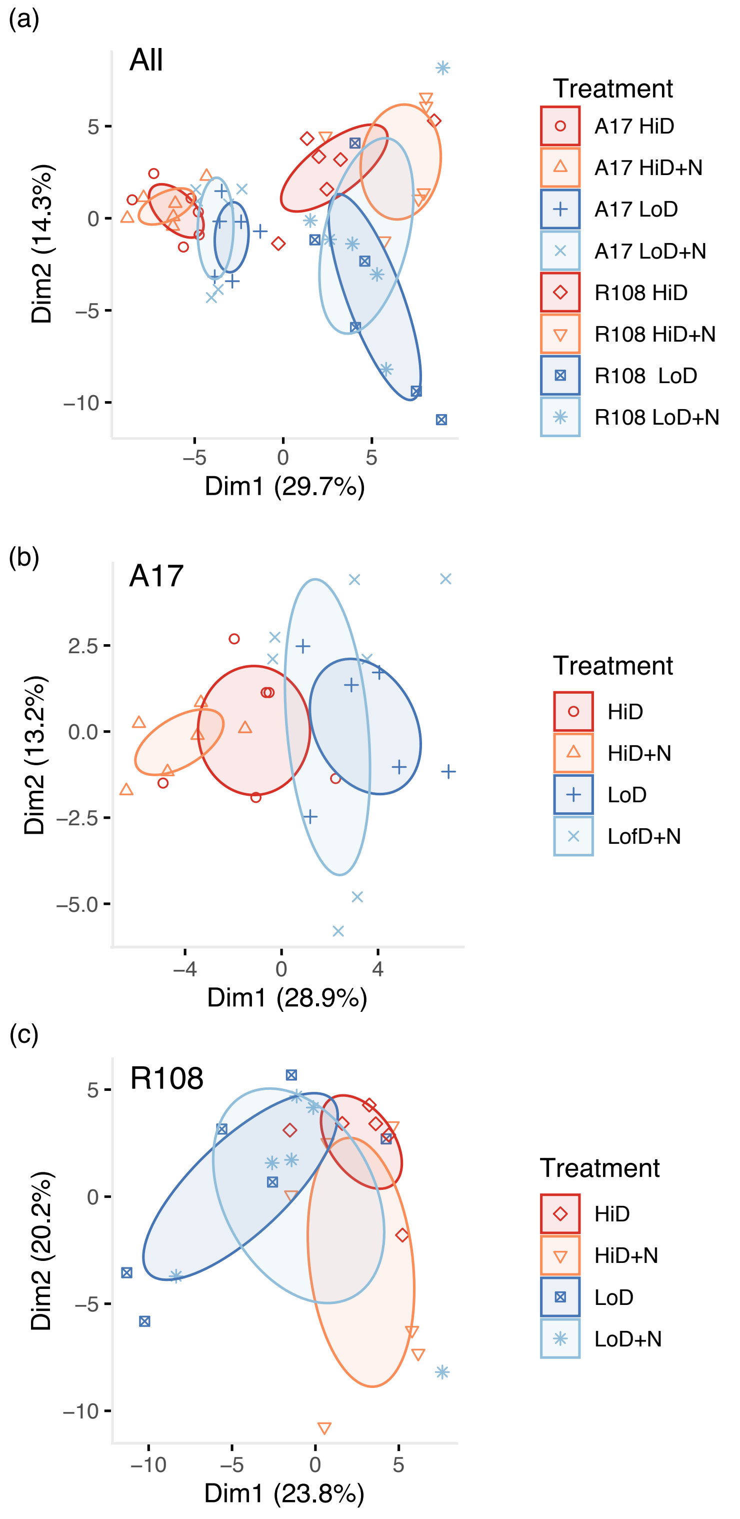

Supplemental Figure S5: Relative frequency of each strain in pairwise and more complex communities A) Pairwise communities of the three focal strains B) C3 strains in C8 and C68 communities, and C) C8 strains in the C68 community and C8 alone.

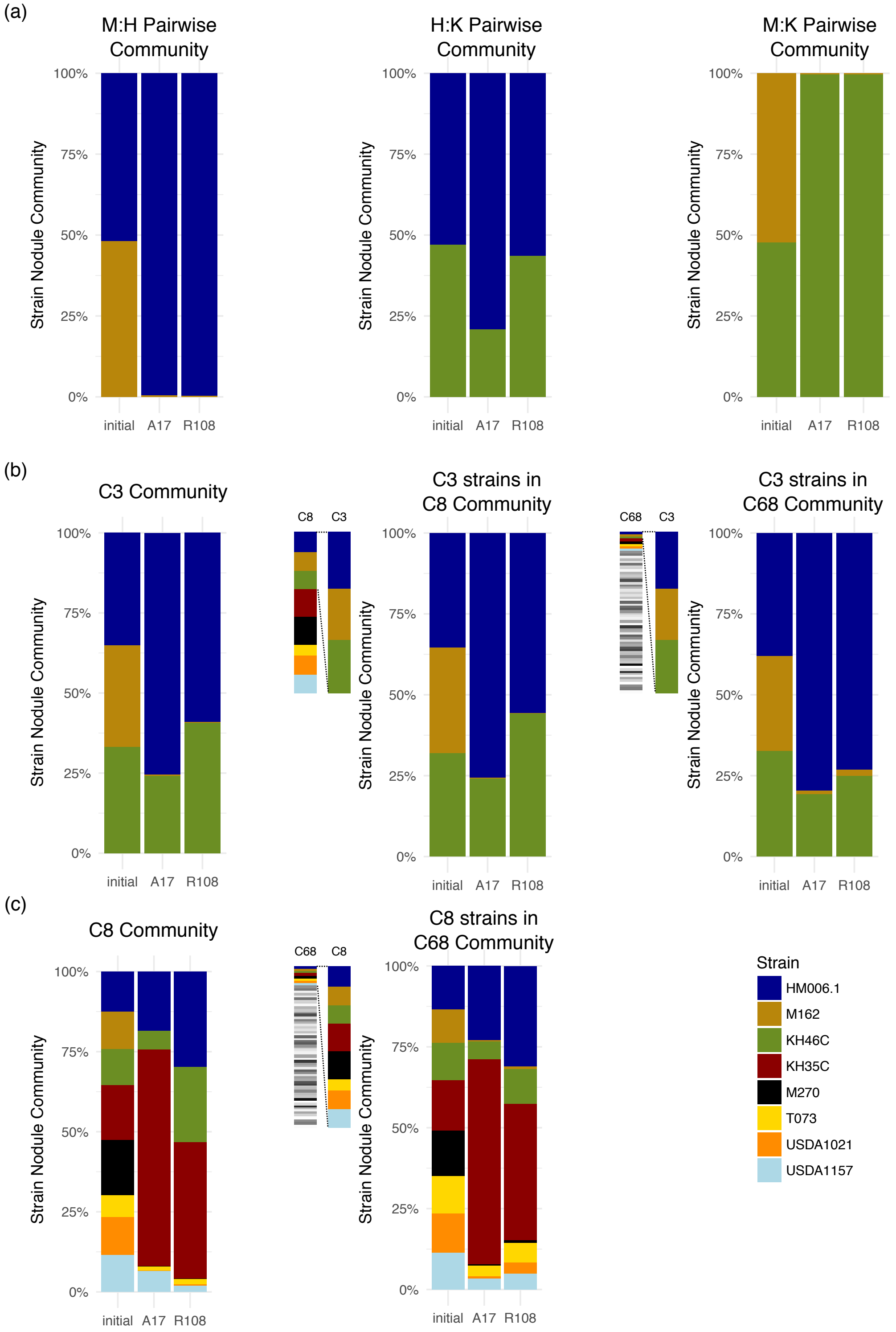

Supplemental Figure S6: Relative frequency of each strain in each replicate in pairwise (a-c) and more complex communities three strain (d) and eight strain communities (e).

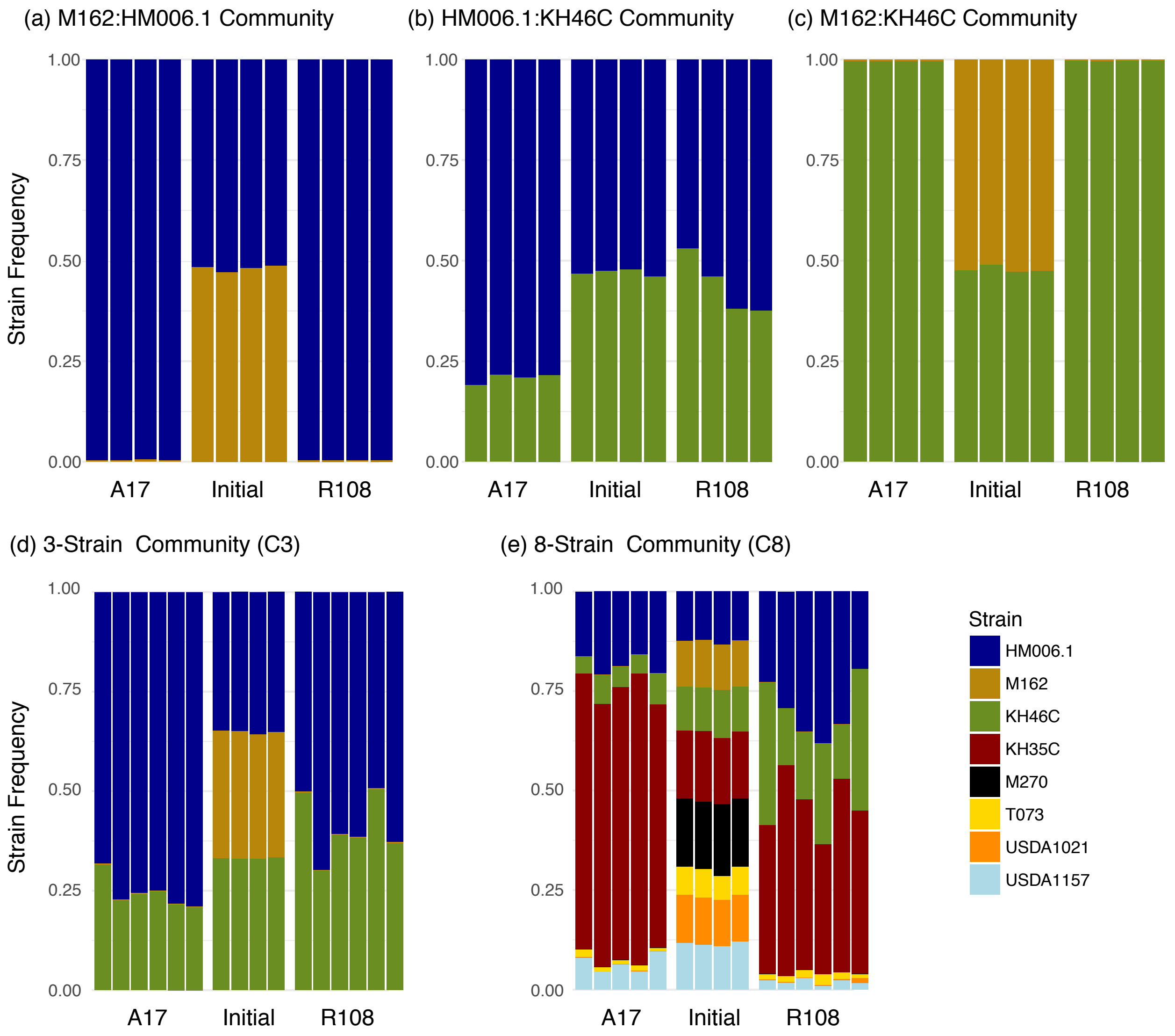

Supplemental Figure S7: Plant biomass traits for Nitrogen x Density experiment including root: shoot ratio (a), dry root biomass (b), and dry shoot biomass (c). ANOVA model results can be found in Supplemental Table S3.

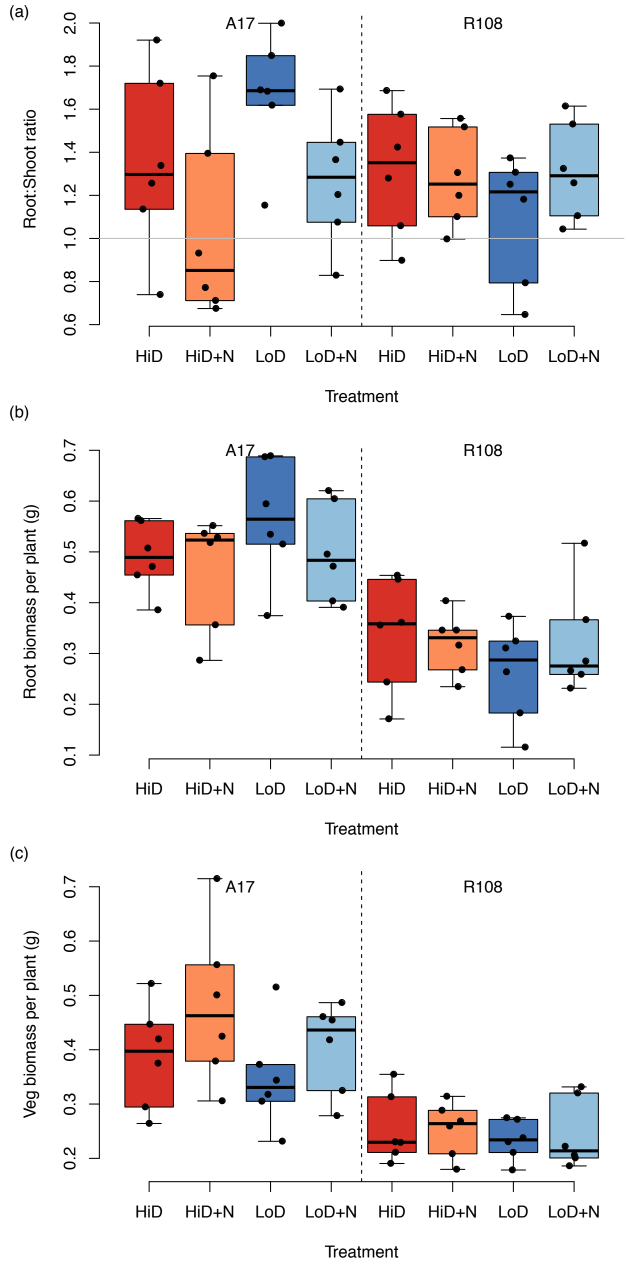

Supplemental Figure S8: Plant phenotypes when A17 and R108 hosts were grown with increasingly complex strain communities: 68 strains (C68), eight strains (C8), three strains (C3), and two strains (H:K, M:H, and M:K). Statistical tests on all six traits can be found in Supplemental Table S4.

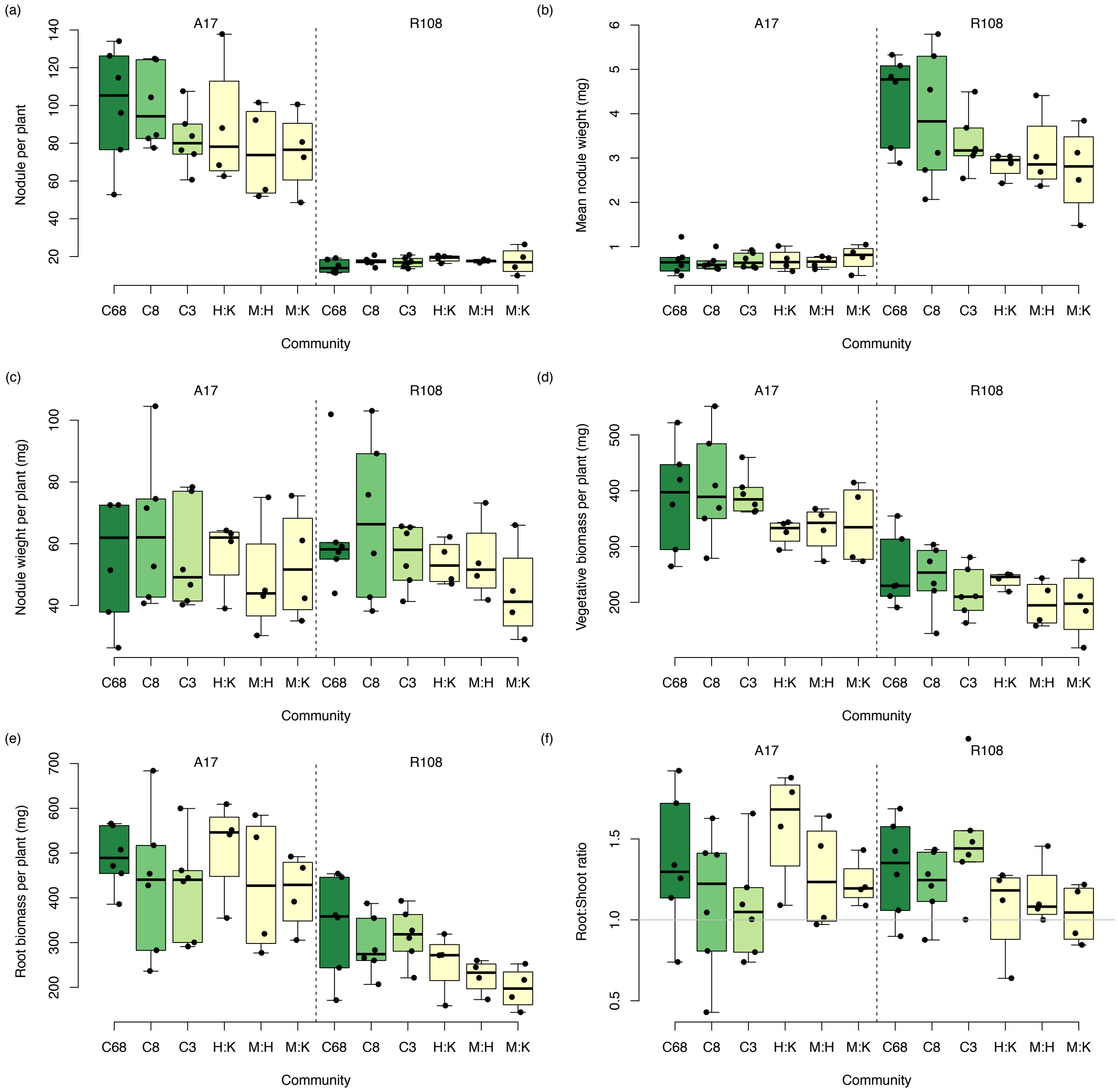

Supplemental Figure S9: Nodule numbers in sequenced pool sizes for (a) the Nitrogen by Density experiment (all plants inoculated with the C68 community) and (b) plants from the community complexity experiment. For R108, nodules were harvested and pooled from all plants in a pot, but due to the large numbers of nodules on A17 only nodules from ~ half of A17 plants were harvested per pot.

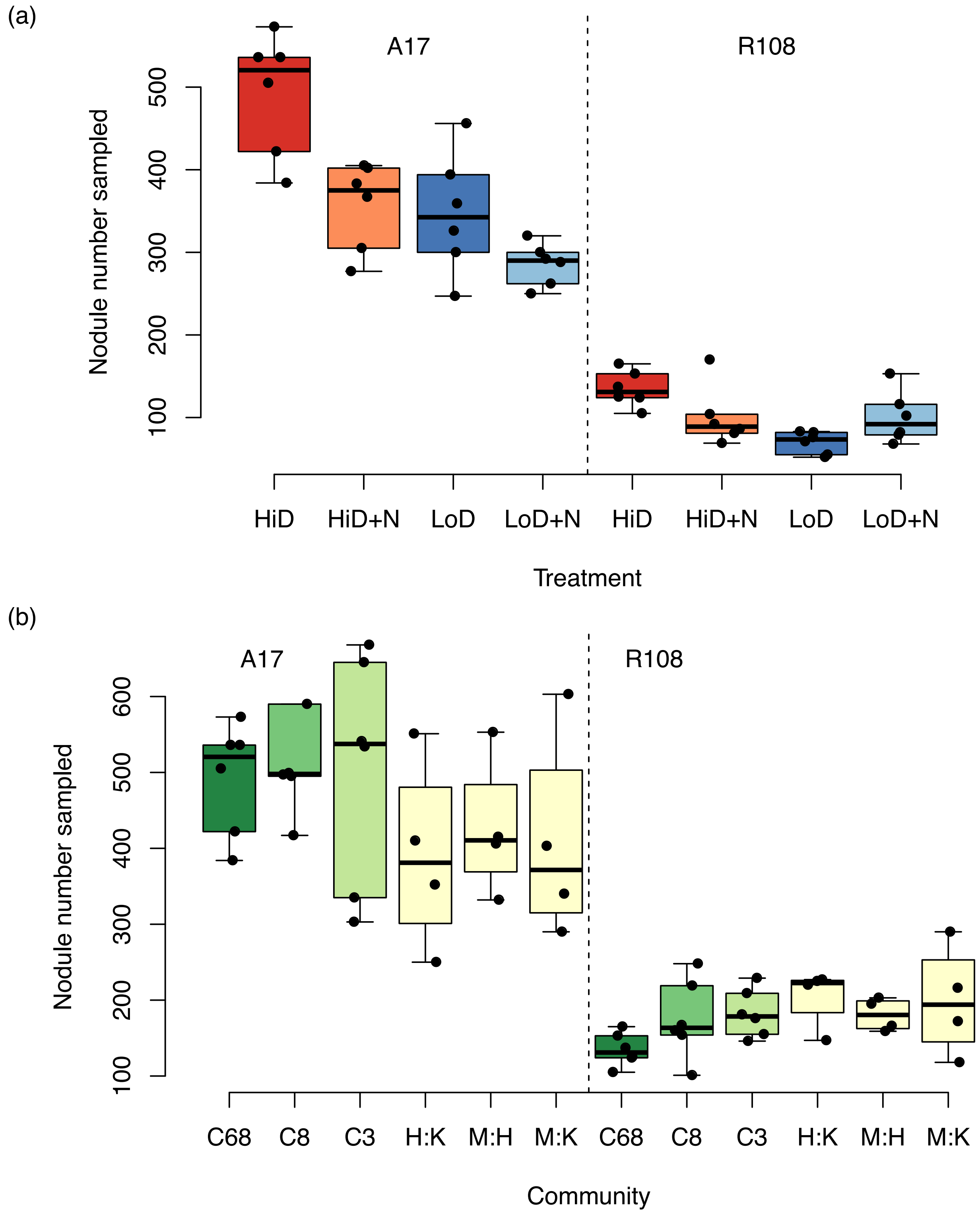
